# Unique protein features of SARS-CoV-2 relative to other *Sarbecoviruses*

**DOI:** 10.1101/2021.04.06.438675

**Authors:** Matthew Cotten, David L. Robertson, My V.T. Phan

**Affiliations:** MRC/UVRI & LSHTM Uganda Research Unit, Entebbe, Uganda; MRC-University of Glasgow Centre for Virus Research, Glasgow, UK

**Keywords:** SARS-CoV-2, proteome changes, *Sarbecovirus* evolution, spike protein changes

## Abstract

Defining the unique properties of SARS-CoV-2 protein sequences, has potential to explain the range of Coronavirus Disease 2019 (COVID-19) severity. To achieve this we compared proteins encoded by all *Sarbecoviruses* using profile Hidden Markov Model similarities to identify protein features unique to SARS-CoV-2. Consistent with previous reports, a small set of bat and pangolin-derived *Sarbecoviruses* show the greatest similarity to SARS-CoV-2 but unlikely to be the direct source of SARS-CoV-2. Three proteins (nsp3, spike and orf9) showed differing regions between the bat *Sarbecoviruses* and SARS-CoV-2 and indicate virus protein features that might have evolved to support human infection and/or transmission. Spike analysis identified all regions of the protein that have tolerated change and revealed that the current SARS-CoV-2 variants of concern (VOCs) have sampled only a fraction (~31%) of the possible spike domain changes which have occurred historically in *Sarbecovirus* evolution. This result emphasises the evolvability of these coronaviruses and potential for further change in virus replication and transmission properties over the coming years.

## Introduction

Since the first report of Coronavirus Disease 2019 (COVID-19) caused by SARS-CoV-2 in December 2019 in Wuhan city, China (Li et al. 2020)(Yang et al. 2020) and the World Health Organisation declaring COVID-19 a global pandemic in March 2020, the disease has proceeded to affect every part of the world. The SARS-CoV-2 virus belongs to the *Coronaviridae* family of enveloped positive-sense single-stranded RNA viruses, *Betacoronavirus* genus, *Sarbecovirus* subgenus. Other *Sarbecoviruses* include SARS-CoV (the coronavirus causing the SARS outbreak in 2002-2004) and a large number of SARS-like bat viruses. The genomes of *Sarbecoviruses* are 30kb in length, encoding >14 open reading frames (ORFs). Among the structural proteins, the spike protein plays a crucial role in virus host-cell tropism, host range, cell entry and infectivity, and is considered the main protein target for protective immune responses. Other virus ORFs encode structural and accessory proteins, many of which modulate important host responses to infection.

Investigation of the evolutionary history of SARS-CoV-2 shows a clear link to *Sarbecoviruses* circulating in horseshoe bats although no direct animal precursor for SARS-CoV-2 has been identified (Andersen et al. 2020) (Boni et al. 2020) (Zhang et al. 2020) (H. Zhou et al. 2020) (Lam et al. 2020). We sought to identify unique peptide regions of SARS-CoV-2 compared to all available *Sarbecoviruses* to determine viral features that might be unique to SARS-CoV-2 and that might have allow the virus to infect, replicate and transmit efficiently in humans. Such a comparative analysis of viral proteins might provide insights into the origin of the virus and identify the conditions that led to the zoonosis to humans, efficient spread without the need for much, if any, adaptation (MacLean et al. 2021), as well as providing leads for drug and immune targets for effective treatments.

## Results and Discussion

### Protein domains and profile hidden Markov models

We have explored the genomes across the *Sarbecovirus* subgenus using profile hidden Markov models (pHMMs). pHMMs can provide a detailed statistical description of an amino acid sequence and can be used to detect related domains found and to document their differences from a reference domain (Eddy 1998) (Eddy 1996). Efficient tools for preparing and comparing pHMMs are available in the HMMER-3 package (Eddy 2011). This approach is particularly useful for comparing large or evolutionary divergent genomes. We have recently used these methods to identify and classify diverse coronaviruses in the *Coronaviridae* family (Phan et al. 2018) and to explore large and unwieldy genomes such as those from the African Swine Fever Virus (Masembe et al. 2020). Here pHMMs were used to explore the relationship between SARS-CoV-2 and the other known *Sarbecoviruses* to gain understanding of their evolutionary history and to identify regions of encoded viral proteins that are with static to change or are altered across the *Sarbecoviruses*.

### Genome scans using custom pHMM domains

First, some background on the strategy used here. We sought to define the distance of any query virus genome from the early SARS-CoV-2 genome that first began to infect humans in December 2019. To give two levels of resolution, we generated overlapping 44 or 15 amino acid (aa) peptides from all early lineage B SARS-CoV-2 encoded proteins then prepared pHMMs using HMMER-3 (see Figure 1a). The resulting libraries of pHMMs were then used to survey domain diversity across query coronaviruses relative to the initial 2019 SARS-CoV-2. For each pHMM match to a related sequence, a bit-score is generated which provides a metric for how close the query sequence is to the pHMM (Figure 1b). These bit-scores, when collected across an entire viral genome, can provide a sensitive description of similarities and difference between a query genome and the reference genome (Figure 1c). For additional background on the method, Supplementary Figure 1 demonstrates the the sensitivity of pHMMs to detect and distinguish single amino acid substitutions and Supplementary Figure 2 demonstrates the use of pHMMs to identify single amino acid substitution in a crucial region of the SARS-CoV-2 spike protein.

**Figure 1.**
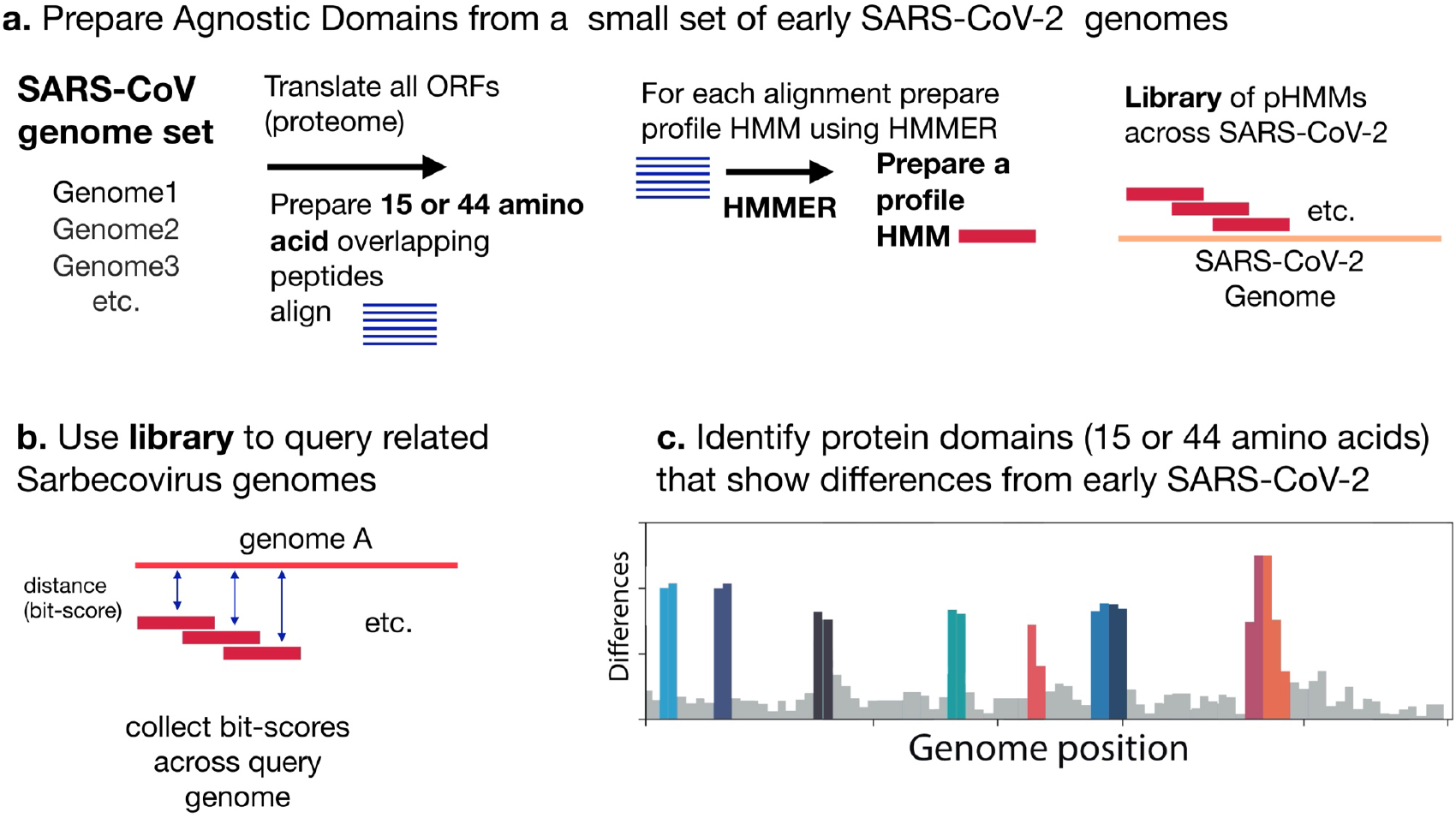
Analysis scheme. **(a)** Profile Hidden Markov Model (pHMM) domains were generated from a set of 35 35 early (Pango) lineage B SARS-CoV-2 genome sequences. All open reading frames were translated and then sliced into either 44 amino acid peptides with a step size of 22 amino acids or 15 amino acid peptides with a step size of 8 amino acid. The peptides were clustered using Uclust (Edgar 2010), aligned with MAFFT (Katoh and Standley 2013) and then each alignment was built into a pHMM using HMMER-3 (Eddy 2011). **(b)** The set of pHMMs were used to query *Sarbecovirus* genome sequences, bit scores were collected as a measure of similarity between each pHMM and the query sequence. **(b)** Bit-scores were gathered and analyzed to detect regions that differ between early SARS-CoV-2 genomes and query genomes.

An initial triage was performed using all available Betacoronavirus genomes from GenBank. All full genomes with the taxon id 694002 were retrieved, genomes with gaps were removed to yield a set of 1480 Betacoronavirus genomes. SARS-CoV-2 genomes were initially excluded from the retrieval and then a set of 27 early lineage B genomes were added as a reference. An additional 5 recently reported bat CoV genomes not yet in GenBank (see Supplemental Table 1) were also added. The 44 aa pHMM library was used to query the Betacoronavirus set. For each genome, the bit-score each of 384 pHMMs from early lineage B SARS-CoV-2 sequences was collected and hierarchical clustering based on the normalized domain bits-scores was performed (Figure 2a). Scores colored with dark to light grey indicating domains identical or close to the corresponding domain from early lineage B SARS-CoV-2 and yellow to orange to red indicating increasing distance. Within the Betacoronavirus set the genomes clustered by their taxonomic group and clusters of OC43, MERS-CoV and SARS-CoV and SARS-CoV-2 were observed. The central region of the *Sarbecovirus* genome is conserved across the genome set with all domains marked as dark or light grey in the Figure 2a This is not unexpected as this central region encodes the viral polymerase, other enzymes and non-surface exposed structural proteins of the virus, which are functionally constrained and less likely to allow change than other regions of the virus. In contrast, the domains displayed in yellow, orange and red in Figure 2a indicated more increasingly divergent regions between early SARS-CoV-2 and the query *Sarbecovirus* genomes (much lower normalized bit scores).

**Figure 2.**
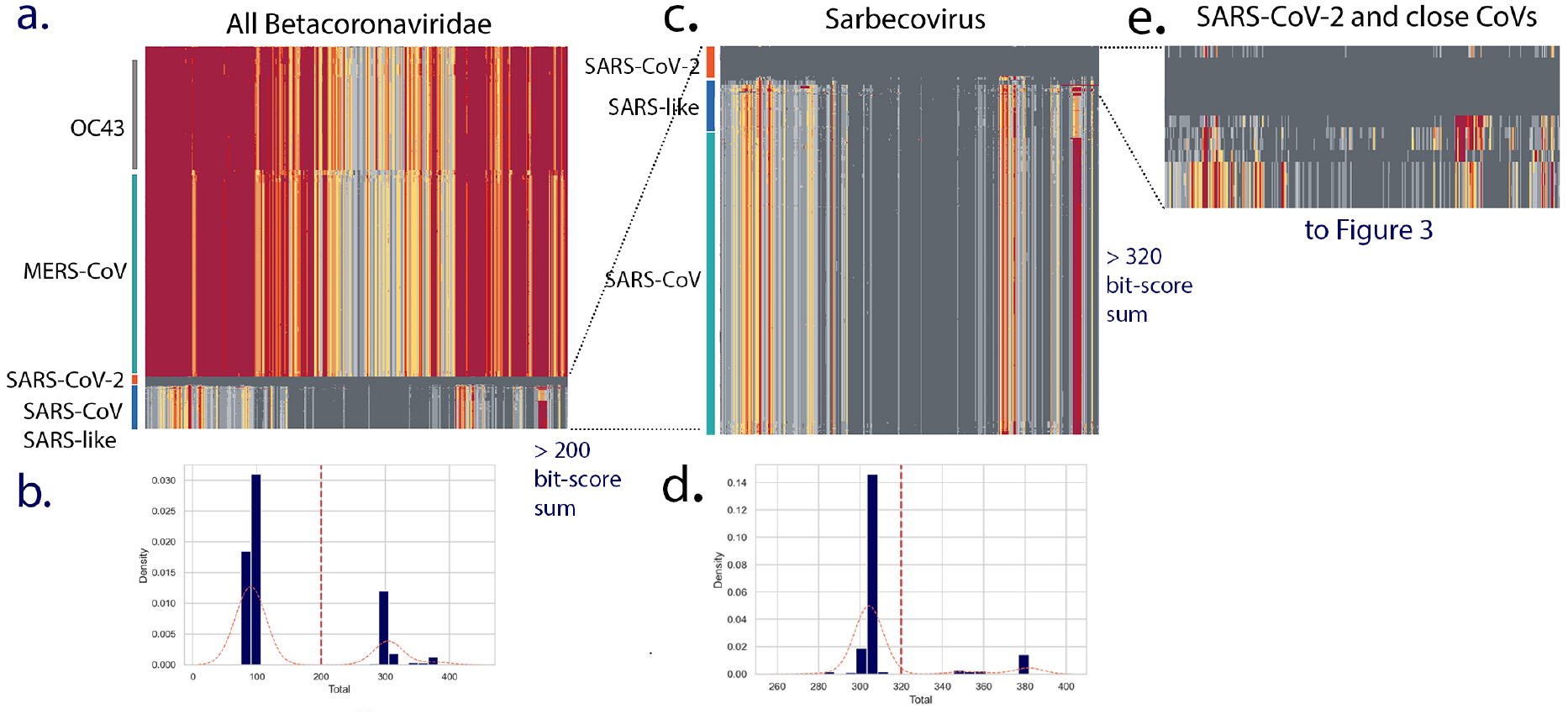
Triage of Betacoronaviruses. All Betacoronavirus genomes (excluding SARS-CoV-2) were retrieved from GenBank using the query ((txid694002[Organism] AND 24000[SLEN]:40000[SLEN] NOT patent)) NOT txid2697049[Organism]) generating a set of 1581 genomes that were screened to remove genomes with gaps to yield a set of 1480 genomes. A small set of 27 early lineage B SARS-CoV-2 genomes from December 2019/January 2020 were added as markers. A library of 44 amino acid pHMMs prepared from early B SARS-CoV-2 genomes was used and the bit-scores for each pHMM were gathered and used to cluster the genomes **(a)** with each domain score indicated by color (dark grey = 1= very similar to SARS-CoV-2 to red = low = distant from SARS-CoV-2). The total bit-scores sum for each genome was calculated (see histograms of all total bitscore sums **(b))**. A total bitscore sum of 200 was used to select for the CoV genomes most similar to SARS-CoV-2. The clustering of this subset of CoV genomes **(c)** included the SARS-CoV-2 genomes, a large number of SARS-CoV genomes and a smaller number of SARS-like CoVs. A cut off of 320 for total bit-scores sum **(d)** was used to identify the closest CoV genomes which were then used for the subsequent analyses reported in Figures 3, 4 and 5.

The sum of the entire set of bit-scores for a genome was then used to calculate a distance from the early SARS-CoV-2 genome. A histogram of these bit-scores sums show several peaks (Figure 2b) with majority of the Betacoronavirus genomes (mostly OC43 and MERS-CoV) clustering around 100 units and a subset of virus genomes with bit-scores >200 units. This >200 set included the SARS-CoV-2, SARS-like CoVs from bats and all the SARS-CoV genomes (Figure 2c). A second triage retained a set of close genomes all with bit-score sums >320 (Figure 2e) that was used more detailed analysis. For simplicity, the 27 early B SARS-CoV-2 genomes in the set (which were nearly identical) were reduced to 5 resulting in a 19 genomes in the close set: 14 bat/pangolin CoV and 5 SARS-CoV-2 (Figure 2e).

We next focused on the bat *Sarbecoviruses* with closest similarity to SARS-CoV-2 in at least part of their genomes due to recombinant histories (see Supplementary Table 1 for genome details and references). The clustermap and variance analysis (Figure 3a) showed higher similarity across most of the genome (dark grey sectors) with three proteins (nsp3, spike and orf9) displaying reduced bit-scores compared to SARS-CoV-2 (Figure 3a, yellow, red domains). These differing regions between the bat *Sarbecoviruses* and SARS-CoV-2 indicate virus protein features that might have evolved to support human infection and/or transmission. The spike differences are explored in detail below however it may be important to consider nsp3, ORF9 (and perhaps nsp4 and ORF8) in future analyses.

**Figure 3.**
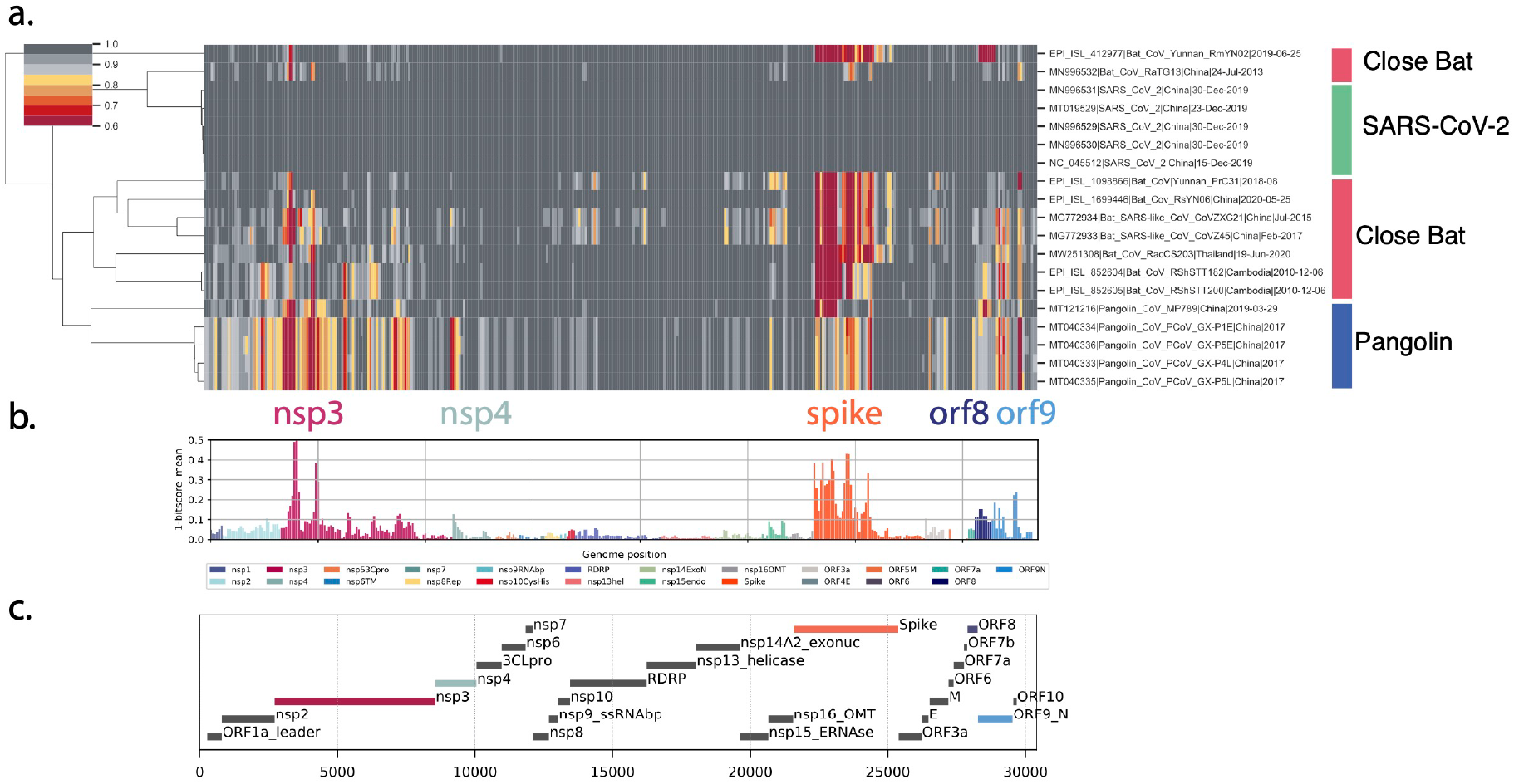
Proteome differences in SARS-CoV-2 versus close bat *Sarbecoviruses*. All forward open reading frames from the 35 early lineage B SARS-CoV-2 genomes were translated, and processed into 44 aa peptides (with 22 aa overlap), clustered at 0.65 identity using Uclust (11), aligned with MAAFT (12) and converted into pHMMs using HMMER-3 (Eddy 2011). The presence of these domains was sought in a set of *Sarbecovirus* genomes plus the SARS-CoV-2 genomes. These were then clustered using hierarchical clustering based on the normalized domain bit-scores (e.g. the similarity of the identified query domain to the reference lineage B SARS-CoV-2 domain). Each row represents a genome, each column represents a domain. Domains are displayed in their order across the SARS-CoV-2 genome, Red = low normalized domain bit-score (lower similarity to lineage B SARS-CoV-2), i.e., higher distance from lineage B SARS-CoV-2, darkest grey = normalized domain bit-score = 1, i.e., highly similar to lineage B SARS-CoV-2. Groups of coronaviruses are indicated to the right of the figure. **(a)** Domain differences across the *Sarbecovirus* subgenus. **(b)** For each domain the mean bit-score was calculated across the entire set of *Sarbecovirus genomes* and the value 1-mean bit-score was plotted for each domain. Domains are coloured by the proteins from which they were derived with the colour code indicated below the figure. **(c)** Schematic of open reading frames or protein products of SARS-CoV-2.

### Spike changes with 15 amino acid domains

Using the same strategy described in Figure 3, we performed a triage of the Betacoronaviruses with 15 amino acid pHMMs prepared from early lineage B spike protein (Figure 4) and selected for CoV genomes encoding close Spike proteins. The high bitscore spike set largely overlapped with the high bit-score full genome set suggesting that spike is a good surrogate for full genome homology.

**Figure 4.**
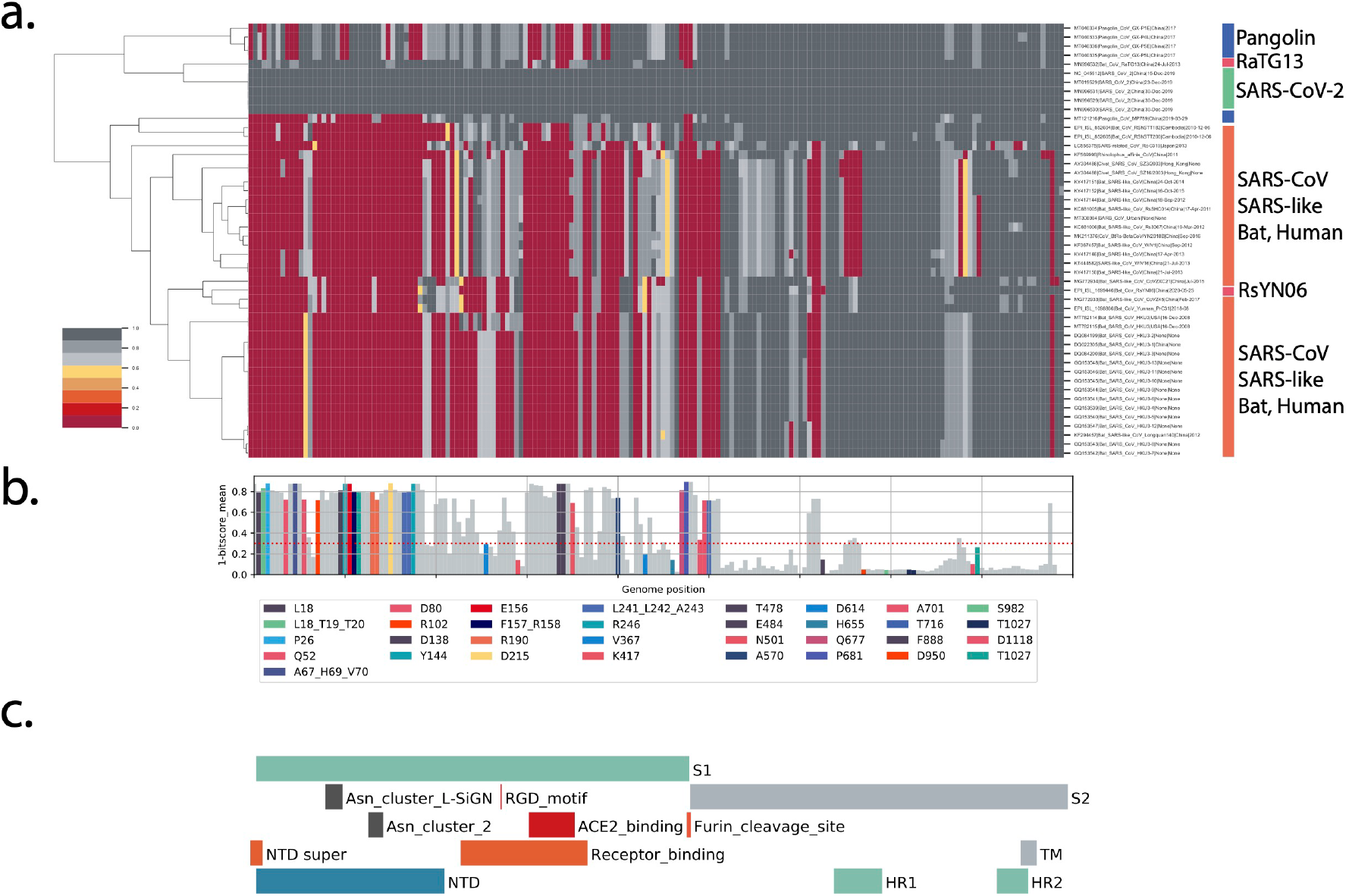
Spike differences in SARS-CoV-2 versus close bat *Sarbecoviruses*. All forward spike open reading frames from the 35 early lineage B SARS-CoV-2 genomes were translated, and processed into 15 aa peptides (with 8 aa overlap) and processed into an pHMM library as described in Figure 2. **(a)** Shows a hierarchical clustering of 15 amino acid domain bit-scores. **(b)** Shows the 1-mean of each domain bit-scores across the genome set, domain values, individual domains that span known amino acid changes in the 6 VOC and VOIs (B.1.1.7, B.1.351, B.1.525, P.1, B.1.617.2 and A.23.1) are colored (see key below panel B). **(c)** The locations of important spike protein features are indicated. NTD: N-terminal domain, RBD: receptor-binding domain, S1: spike 1, S1: Spike 2, TM: transmembrane domain, HR1: helical repeat 1, HR2: helical repeat 2, NTD super: N-terminal domain supersite.

Key features of the spike protein are outlined in Figure 4c. The analysis revealed regions of spike that historically have tolerated change. In general the S1 subunit of spike (the amino-terminal half of the protein) displayed a large amount of diversity with most of the low score domains (more distant from SARS-CoV-2, marked in red) concentrated here (Figure 4a). This is consistent with the surface exposure of S1 on the virion and with protein changes driven by pressure to avoid immune responses. The central ACE2 receptor binding region (Figure 4c) was very different between the close *Sarbecoviruses* and SARS-CoV-2. The furin cleavage site at the junction between the S1 and S2 domains (Figure 4c) is also a region showing a lot of diversity across the *Sarbecovirus* spikes (Figure 4a) and appears completely unique to SARS-CoV-2. This has been discussed in detail (Hoffmann et al. 2020) and is also a site of frequent change in the current SARS-CoV-2 Variant of Concern (VOC) spike sequences with Q677, P681 and T717 flanking the furin site showing changes (Figure 4c).

The domains that where amino acid changes have appeared in VOC spike proteins are marked in color (Figure 4b) and largely appear in domains with high variation (Figure 4b, 1-mean bit-score > 0.3) suggesting that *Sarbecoviruses* have made changes in these regions in previous evolutionary periods and are continuing to change in SARS-CoV-2 evolution. Of the 88 spike domains showing high variation in *Sarbecoviruses* (1 - mean bit-scores >= 0.3 units), only 27 of the domains (31%) have accumulated substitutions or deletions. This indicates a very large potential in the SARS-CoV-2 spike protein for tolerating future change. Important regions that have shown high levels of historical change are the NTD, the RBD and the furin cleavage site and flanking regions.

### Global proteome similarities

As described in Figure 2, a measure of the total protein distance between the SARS-CoV-2 and any query *Sarbecovirus* can be obtained by summing the normalized bit-scores (SNBS) across the entire query proteome. We examined SNBSs grouped by virus host for the 44 amino acid total genome analysis and the 15 amino acid spike gene analysis. The potential role of pangolins as an amplifying intermediate host of SARS-CoV-2 is important to document securely, to guide efforts to prevent or prepare for future zoonotic events. A small number of *Sarbecoviruses* have been identified in samples from trafficked pangolins in China (Liu et al. 2019) (Lam et al. 2020) (Xiao et al. 2020), yet there is no direct evidence that pangolins host the virus in their natural environment. It is thus likely these pangolins identified in China were infected by viruses encountered after transport to China, consistent with reports of disease in these animals. Five CoV sequences from pangolins were included in this analysis (Supplementary Table 1), including four generated by Lam *et al*. (Lam et al. 2020) after sequencing the original samples described by Liu *et al*.(Liu et al. 2019); a 5th genome (MP789) was deposited by Liu et al..

The bat coronavirus genome RaTG13 (GenBank MN996532.1) was identified as closely related to the SARS-CoV-2 lineage (P. Zhou et al. 2020) and supports a bat coronavirus being the zoonotic source of the epidemic, although despite the close genetic distance it is too far in time (decades) for RaTG13 itself to be a direct source of the pandemic SARS-CoV-2 virus (Boni et al. 2020). The next closest bat coronavirus RsYN06, shows some regions of even close identity to SARS-CoV-2 than RaTG13 (H. Zhou et al. 2020) (Figure 4a) due its possible recombinant nature. A single pangolin derived SARS-CoV-2 (MP789) showed an SNBS value that was also elevated but not as high as the RaTG13 (Figure 5a), the 15 aa spike analysis showed similar patterns except that only the RaTG13 spike displayed the high similarity to SARS-CoV-2 (Figure 5b).

**Figure 5.**
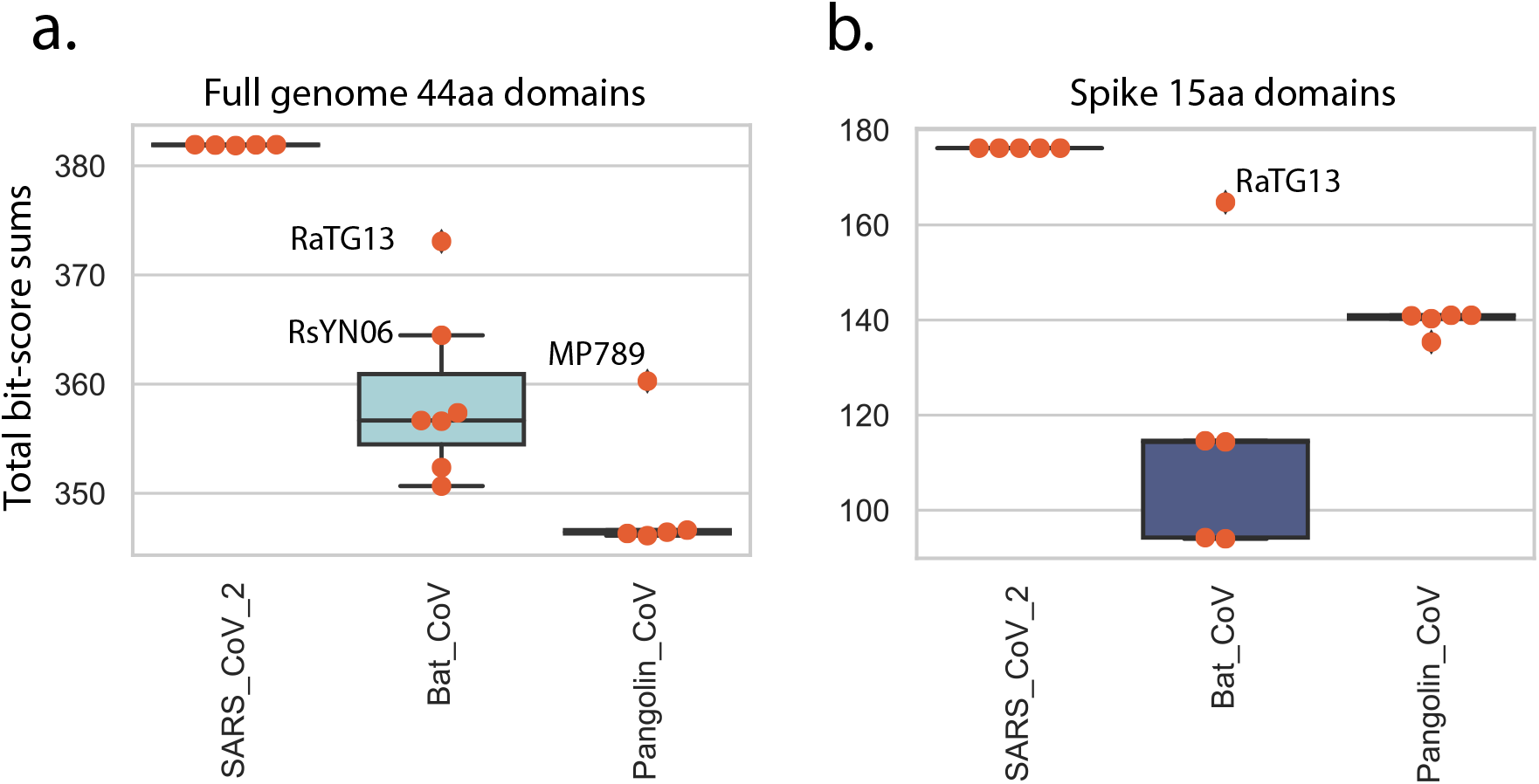
Total domain distances between virus groups. Normalized bit-score sums (NBSS) grouped into SARS-CoV-2 and *Sarbecoviruses* from pangolin, bat, for all domains for each genome were summed. The boxplot shows individual values marked in orange, median values indicated by horizontal black lines, 1st interquartile ranges marked with a box. The identities of several high scoring bat and pangolin genomes are indicated. **(A)** NBSS for 44 aa domains across the entire coronavirus genome. **(B)** NBSS for 15 aa domains across the spike protein.

## Conclusions

What is special about SARS-CoV-2? Spike changes in SARS-CoV-2 compared to the close set of *Sarbecovirus* genomes indicate that the immediate zoonotic source of SARS-CoV-2 is yet to be identified due to the unique nature of the SARS-CoV-2 genome. The more detailed analysis of spike regions in SARS-CoV-2 genomes (Figure 3) revealed the extent of the changes that have occurred across the *Sarbecovirues*. Combined with the current VOC spike changes (from lineages B.1.1.7, B.1.351, B.1.525, P.1, B.1.617.2 and A.23.1), the patterns suggest that SARS-CoV-2 has a great deal of possibilities for further evolution, presumably enabling persistence and avoid immune responses. This emphasises the importance of genomic variant surveillance for monitoring for further changes in virus biology that may have implications for spread and disease severity. Vaccine producers should be prepared to accommodate such spike changes in the next generation of vaccine updates. In addition to the spike protein, additional regions of high variance were observed in the nsp3 across all *Sarbecoviruses* (Figure 2) in close bat and pangolins (Figure 3).

The high variance regions flanked and partially overlapped the Macro domain, which is frequently associated with ADP-deribosylase activity (Frick et al. 2020)’(Lei et al. 2018). Variance observed in the ORF8 changes across the set was due to frequent deletion of this ORF, suggesting that the encoded protein may be dispensable for human infection. Similar loss of ORF8 was observed with the original SARS-CoV (Chiu et al. 2005) (Tang et al. 2006) and has been observed in several SARS-CoV-2 lineages as the virus adapted to humans (Su et al. 2020) (Gong et al. 2020) (Young et al. 2020). The ORF9 (N protein) variance observed across *Sarbecoviruses* and the changes in this protein in VOC strains suggest an additional region that may be adapting to human replication. The regions of variance identified here may indicate either functional changes in SARS-CoV-2 proteins or amino acid positions that can be changed without impairing the necessary functions of the protein. The relatively high mutation rate of SARS-CoV-2 combined with the unprecedented number of SARS-CoV-2 infections in the world is resulting in massive viral adaptation. Additional experiments are required to distinguish true functional changes from neutral evolution.

Finally, the detailed spike analysis of Figure 4 revealed 88 15aa spike domains showing high variation while only 27 (31%) have accumulated substitutions or deletions in the current epidemic in VOCs and VOIs indicating a large potential for tolerating future change. It is highly likely that a large number of new SARS-CoV-2 variants with changes in these regions will evolve, compatible with similar levels of virus replication but tolerating significant antigenic change in the coming years, unless global SARS-CoV-2 spread is severely curtailed.

## Acknowledgements

We thank all global SARS-CoV-2 sequencing groups for their open and rapid sharing of sequence data and GISAID for providing an effective platform for making these data available. This work was supported by the Wellcome, DFID - Wellcome Epidemic Preparedness – Coronavirus (AFRICO19, grant agreement number 220977/Z/20/Z) awarded to MC. DLR and MC receive funding from the MRC (MC_UU_1201412), by the UK Medical Research Council (MRC/UKRI) and the UK Department for International Development (DFID) under the MRC/DFID Concordat agreement (grant agreement number NC_PC_19060).

## Supplementary Material

**Supplementary Table 1.**
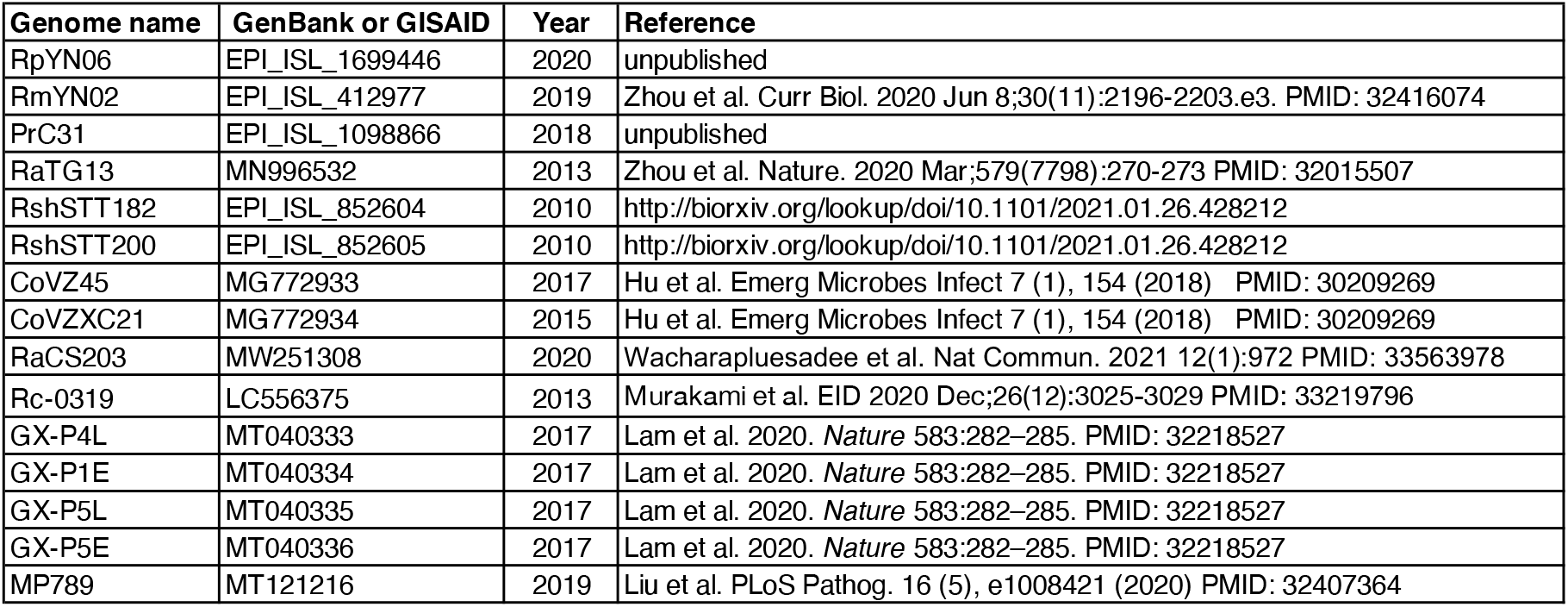
Close bat coronaviruses.

### Supplementary Figure 1 to illustrate pHMM detection of amino acid changes

We sought to illustrate the ability of pHMMs to detect amino acid differences between a reference and a query sequence. A reference peptide containing the twenty amino acids was used to prepare a pHMM. A test set of mutant sequences was prepared by sequentially changing each amino acid to each of the other 20 amino acids. This set of 400 sequences was then queried with the wildtype 20aa profile HMM, the bit-scores describing each match were collection. The distribution of bit-scores from the 400 pHMM matches (Supplementary Figure 1a) was broad, consistent with the methods ability to report not only an amino acid changes but the type of amino acid change. The pattern of all amino acid changes across all 20 AA peptide is displayed in clustermap (Supplementary Figure 1b) with each column corresponding to a single amino acid and each row showing the score if that amino acid were changed to another amino acid. An amino acid that is frequent (e.g. alanine (A)) shows higher bit-scores across the set of changes than rarer amino acids such as cysteine (C), histidine (H), tryptophan (W) or proline (P). This spectrum closely reflects the BLOSUM62 substitution matrix (Henikoff and Henikoff 1992) and demonstrates the capacity of a pHMM match both to detect changes in proteins as they evolve and to distinguish different types of changes.

**Supplementary Figure 1.**
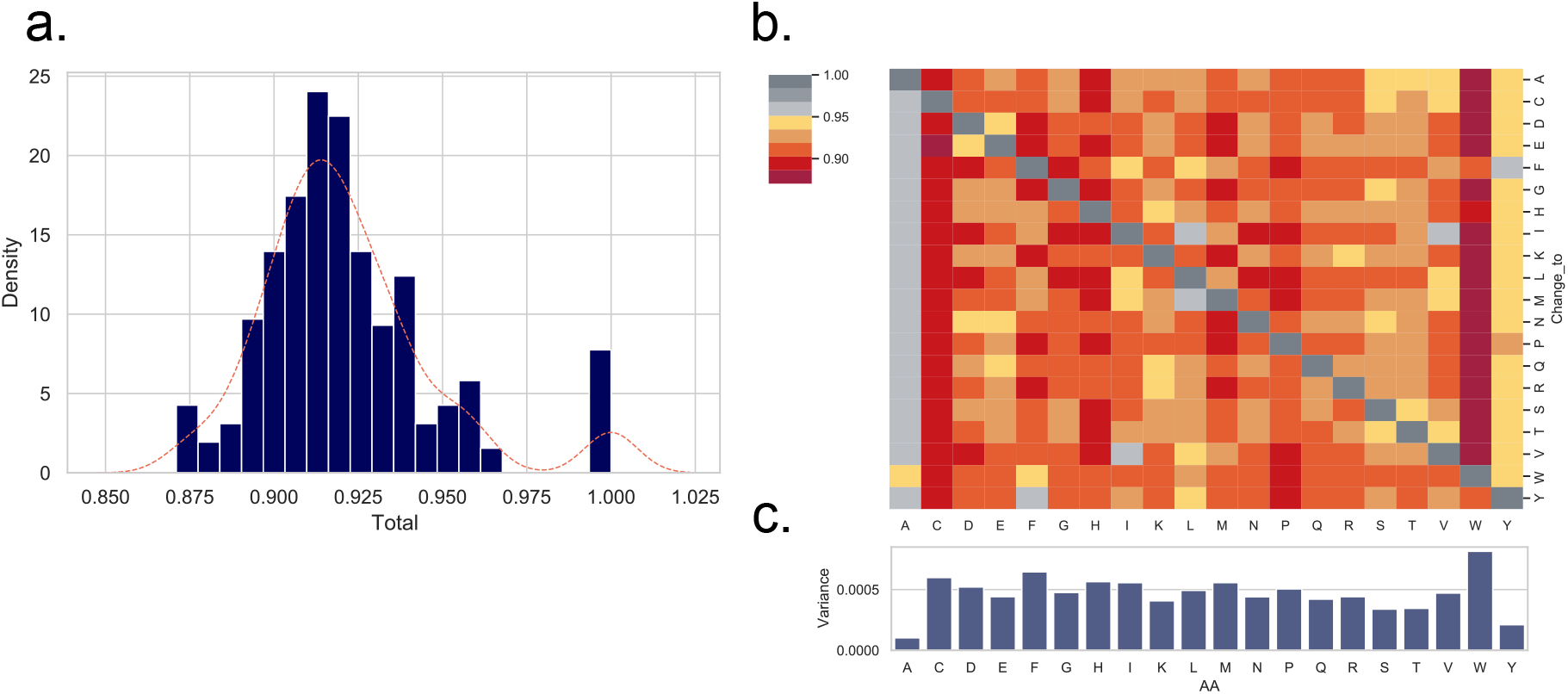
pHMM bit-score values as a measure of the type of amino acid change. A sequence encoding all 20 amino acids (ACDEFGHIKLMNPQRSTVWY) was used to prepare a pHMM. A test set of mutant sequences was prepared by changing each amino acid to each of the other 20 amino acids. This set of 400 sequences was then queried with the wildtype 20aa profile HMM, bit-scores for each match were collected in a matrix. **(a)** a histogram of all observed normalized bit-scores, the peak at 1.00 is due to changes to self (e.g. A to A change). **(b)** heatmap of normalized bit-scores, each columns represents a position in the 20 AA wt peptide, each row represents a change at that position to the indicated amino acid. The normalized bit-scores were color coded with no change from wildtype amino acid (dark grey) to the largest change from the wildtype amino acid (dark red). **(c)** Variance of normalized bit-scores from panel b were calculated for each position.

In a second analysis we examined a peptide sequence spanning the important furin cleavage site in the SARS-CoV-2 spike protein. Mutations in this region have appeared in several VOCs (A.23.1: P681R, B.1.1.7: P681H, B.1.525: Q677H) and we wanted to document the sensitivity of pHMM matching to detect single amino acid changes. Similar to Supplementary Figure 1, we prepared a pHMM from the wildtype 14aa sequence spanning the furin site. For a test set we systematically change position to each of the 20 amino acids and then gathered the bit-scores for the wildtype pHMM matching each test peptide. Similar to Supplementary Figure 1, the range of normalized bit-scores scores included a peak at 1.00 (self sequence matched to self) plus a range of lower values demonstrating the breadth of possible pHMM bits-scores for any possible single amino acid changes in the 14 amino acid peptide (Supplementary Figure 2a). The heatmap of the resulting normalized bit-scores (Supplementary Figure 2b) reveals some patterns. Most changes of the proline adjacent to the cleavage site resulted in a large reduction in bit-scores, whereas other changes resulted in detectable, distinct, but less dramatic bit-scores.

**Supplementary Figure 2.**
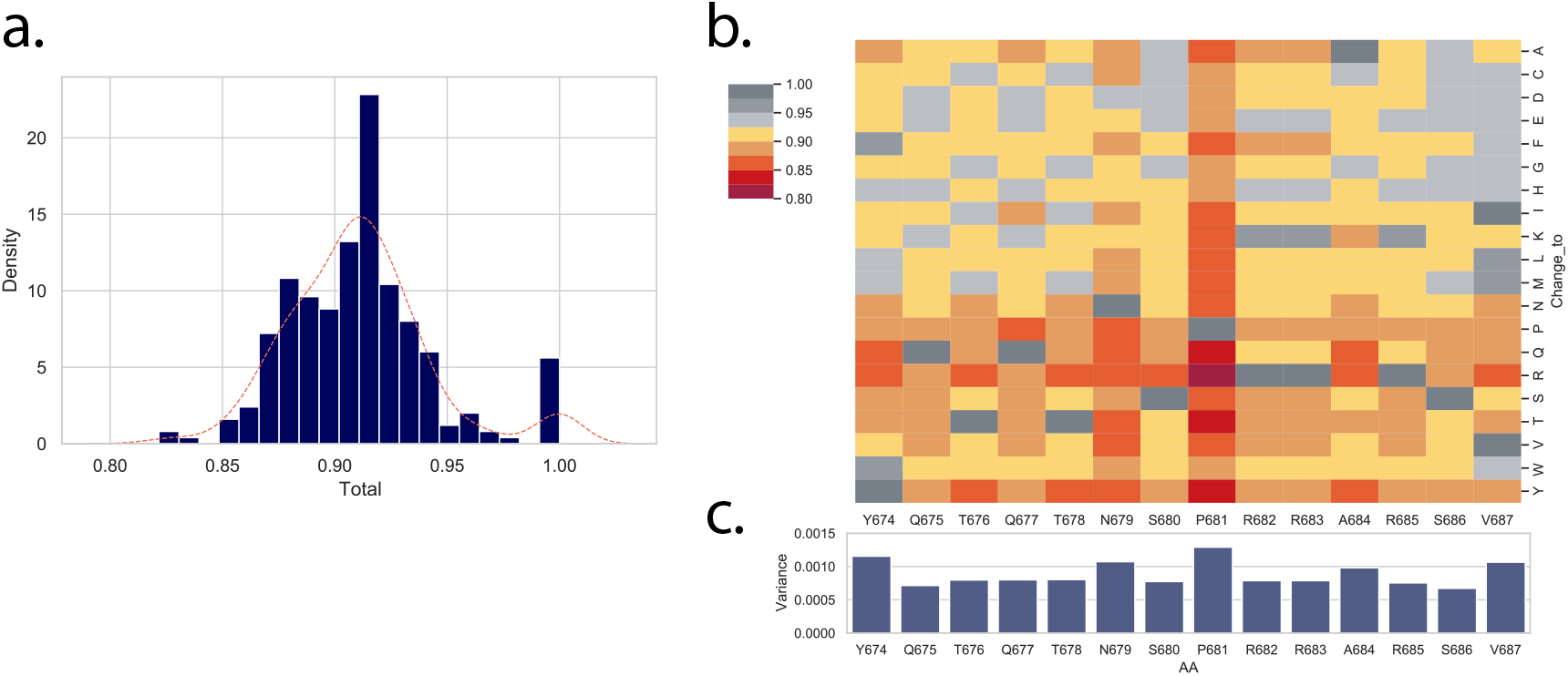
Amino acid changes across the P681 region of the spike protein. A 14 amino acid sequence spanning the SARS-CoV-2 spike position 681 and the adjacent furin cleavage site (YQTQTNSPRRARSV) was used to prepare a pHMM. A test set of mutant sequences was prepared by changing each amino acid to each of the other 20 amino acids. This set of 280 sequences was then queried with the wildtype 14aa pHMM, bit-scores were collected. Panel a, a histogram of all observed normalized bit-scores, the peak at 1.00 due to changes to self (e.g. A to A change). Panel b, heatmap of normalized bit-scores, each columns represents a position in the 14 AA wt peptide, each row represents a change at that position to the indicated amino acid. The normalized bit-scores were color coded with no change from wildtype amino acid (dark grey) to the largest change from the wildtype amino acid (dark red). Panel C. Variance of normalized bit-scores from Panel b were calculated for each amino acid position across the peptide.

